# Recurrence and propagation of past functions through mineral facilitated horizontal gene transfer

**DOI:** 10.1101/2023.01.24.525235

**Authors:** Taru Verma, Saghar Hendiani, Carlota Carbajo Moral, Sandra B. Andersen, Emma Hammarlund, Mette Burmølle, Karina K. Sand

## Abstract

Horizontal gene transfer is one of the most important drivers of bacterial evolution. Transformation by uptake of extracellular DNA is traditionally not considered to be an effective mode of gene acquisition, simply because extracellular DNA is degraded in a matter of days when it is suspended in e.g. seawater. Mineral surfaces are, however, known to preserve DNA in the environment, and sedimentary ancient DNA studies have solidified that there are considerable amounts of fragmented DNA stored in sediments world-wide. Recently the age span of stored DNA was increased to at least 2 Ma. Here, we highlight that fragmented ancient DNA can be fueling the evolution of contemporary bacteria and advocate to consider this route for genetic variation in evolutionary history. We show that *Acinetobacter baylyi* can incorporate 60 bp DNA fragments adsorbed to a wide range of common sedimentary minerals and that the transformation frequencies scale with the mineral surface properties. Further, our results point to interfacial geochemical and sedimentologic processes as facilitators of evolutionary innovation where DNA-molecules are specific to the environment and the processes providing new DNA molecules may also provide the need to evolve. In contrast to heritable stochastic mutations as proposed by Darwin, the access by which bacteria acquire new genomic material at times with increased stress and also needs, would indicate a non-random mechanism that may propel evolution in a non-stochastic manner.

## INTRODUCTION

Horizontal gene transfer (HGT) is recognized as the key source of accelerating genome innovation.^1^ In HGT, extracellular or intracellular DNA from one species can be incorporated into a related or non-related species, and thereby efficiently increase genetic diversity. HGT is considered a common route to adaptation in microbes and has significantly impacted the evolution of life of both prokaryotes and eukaryotes. ^2,3^ HGT through conjugation and transduction is relatively well explored and conjugation is considered a key mechanism for the propagation of antibiotic resistance genes. HGT by transformation, i.e. uptake of extracellular DNA, is currently not considered an effective **mode** of **gene acquisition** because extracellular DNA degrades in a matter of days when it is suspended in seawater or a soil environment. ^4-7^ However, advances made in the field of sedimentary ancient DNA underline that adsorption of DNA to mineral surfaces significantly enhances DNA preservation. ^8^ The time-span of DNA preservation was just increased from 1.45 million years to at least 2 Ma years. Currently, there is about 0.4 Gt of extracellular DNA associated with sediments just in the top 10 cm of the ocean floor ^9^ and in principle, all sedimentary minerals hold a potential to adsorb some amount of extracellular DNA.

The interplay between organic molecules and mineral surfaces can be described by interfacial geochemical principles. The adsorption capacity determines how much DNA the mineral can adsorb at a given DNA concentration and is often influenced by the environmental conditions such as solution pH, salinity, etc. The minerals comprising sedimentary systems are mineralogically diverse with distinct surface characteristics such as types of crystal faces, surface chemistries, surface composition and active site densities -all influencing the associations with biomolecules. Minerals with an overall positive surface charge and high densities of active sites such as clay edges, carbonates and oxyhydroxides are shown to have a higher DNA adsorption capacity and are less dependent on solution composition than negatively charged minerals such as clay basal planes and silicate minerals. The negatively charged minerals depend on positively charged background ions to facilitate binding of the DNA. A recent study combining molecular dynamic simulation and atomic force microscopy (AFM) showed that the higher the charge density on topographic surface features, as well as of solution cations (ionic potential), the more the DNA is immobilized. ^10^ Besides preserving adsorbed DNA, mineral surfaces can also act as vehicles for transport of the DNA. When adsorbed to a mineral surface, DNA representing niches in a given environment can enter a distal ecosystem through common sedimentary pathways, provided the DNA-mineral association can survive a potential change in chemical environment associated with relocation.

The frequency of HGT events is considered low, ^11^ but we argue that the likelihood of a transformation event from mineral adsorbed DNA is enhanced by the anchoring of many bacteria to mineral surfaces and the vast amount of DNA stored in sediments. Bacteria are found to colonize any surface in most sedimentary systems, ^12,13^ and have ample opportunities to encounter mineral surfaces. They associate with mineral surfaces suspended in the water column, and minerals depositing or deposited on e.g. a sea floor. Contemporary bacteria also inhabit sedimentary deposits formed in the past gaining access to past genes. It is generally found that if an acquired gene encodes beneficial traits in a given environment, it will provide the host with a fitness advantage, resulting in stabilization of the trait(s) in the population due to natural selection. ^14^ The mixing of modern microbes and ancient DNA in principle represents a pathway for gene transfer, across a scale of time and space, unrivaled by any other avenue.

Transformation of mineral-adsorbed plasmid and whole genomic DNA was shown from clays (kaolinite and montmorillonite) and iron oxy-hydroxides (goethite). ^15,16^ While DNA molecules are found to be associated with many sediments including soils, ^17^ it is highly unlikely that the DNA will remain intact for long timescales, owing to degradation processes such as spontaneous hydrolysis, action of nucleases and UV-mediated degradation. Common for sedimentary DNA molecules is that they are highly fragmented and degraded and usually shorter than 100 base pairs (bp). ^18^,^19^ In principle, transformation of bacteria by short DNA fragments can occur for any bacterial species, owing to the simple mechanism of DNA uptake and the sheer abundance of fragmented DNA in the environment. ^20^ Integration of small DNA fragments in the bacterial genome is spontaneous and very simple in comparison to natural transformation of kilobase size DNA molecules (RecA-mediated) which requires a strong sequence homology. During uptake of fragmented DNA, the bacteria degrade the DNA double helix and release single-stranded DNA into the cell. When the bacterial genome replicates, the released single stranded fragments can act as primers of new DNA and bind to the open chromosome near the replication fork, thereby incorporating foreign DNA in the genome of one of the daughter cells. ^20^ Successful transformation of bacteria by short extracellular DNA fragments has been shown for *Acinetobacter baylyi*. ^21^ *A. baylyi* is a soil bacterium that is naturally highly competent and recombinogenic with no specific sequence requirements for DNA uptake. Despite the relative ease of incorporation, uptake of fragmented DNA associated with mineral surfaces remains unexplored and its evolutionary and ecological implications are not considered.

The involvement of a surface in a transformation event introduces the interfacial reactions between the molecule, surfaces and/ or bacteria as potential co-drivers for the uptake process. i.e. it is yet not known if a strong DNA-mineral affinity (bond strength, density of binding sites, types of bonds etc) decrease or even prevent uptake by decreasing the availability of the DNA molecule to the microbe. Here, we explore if mineral-adsorbed short DNA fragments are available for bacterial uptake via HGT in *A. baylyi*. We pre-adsorbed 60 bp DNA molecules encoding the *trpE* gene, to a broad range of environmentally relevant soil minerals, (iron oxides and hydroxides (hematite and goethite); clays (kaolinite and mica); carbonates and non-clay silicates (quartz)) and measured the bacterial uptake efficiency per μg of the adsorbed DNA for nutritionally deficient and rich conditions. We evaluate how mineral surface characteristics impact transformation efficiency and further set the results in sedimentologic and ecologic contexts and elaborate on how this process could influence the evolution of life as suggested by Sand and Jelavic. ^22^

## RESULTS AND DISCUSSION

### DNA uptake by transformation

The experimental strategy and protocols used for DNA adsorption and horizontal gene transfer experiments are depicted in Materials and Methods Figure MM-1.

#### DNA adsorption

All minerals used in this study display a range of different crystal faces, surface chemistries, surface composition and active site densities (nm^-1^) **(Table S1.1)**. All these parameters give rise to differences in surface charge, adsorption behavior and adsorption parameters where the main parameters are the density of active sites and the surface charge. Figure 2 shows the distinct adsorption capacities between the minerals for 60 bp DNA fragments encoding *trpE* in 150 mM NaCl solution. Goethite, kaolinite, calcite and mica appeared to follow a Langmuir isotherm and a maximum adsorption capacity could be derived. Hematite and quartz, conformed well to a Freundlich isotherm with no surface saturation within the range of equilibrium DNA concentration tested (R^2^ values for hematite: 0.97 for Freundlich vs 0.91 for Langmuir: R^2^ values for quartz: 0.8 for Freundlich vs 0.1 for Langmuir) (Fig. 1 and Fig. S1.1). Plotted as μg DNA /mg mineral, goethite showed the highest maximum adsorption capacity (∼45 μg/mg), followed by hematite (∼10 μg/mg) and kaolinite (∼5 μg/mg)) (Fig. 1(a)). Calcite, mica, and quartz adsorbed the lowest amount of DNA per mg of mineral (0.1-0.2 μg/mg).

**Figure 1:**
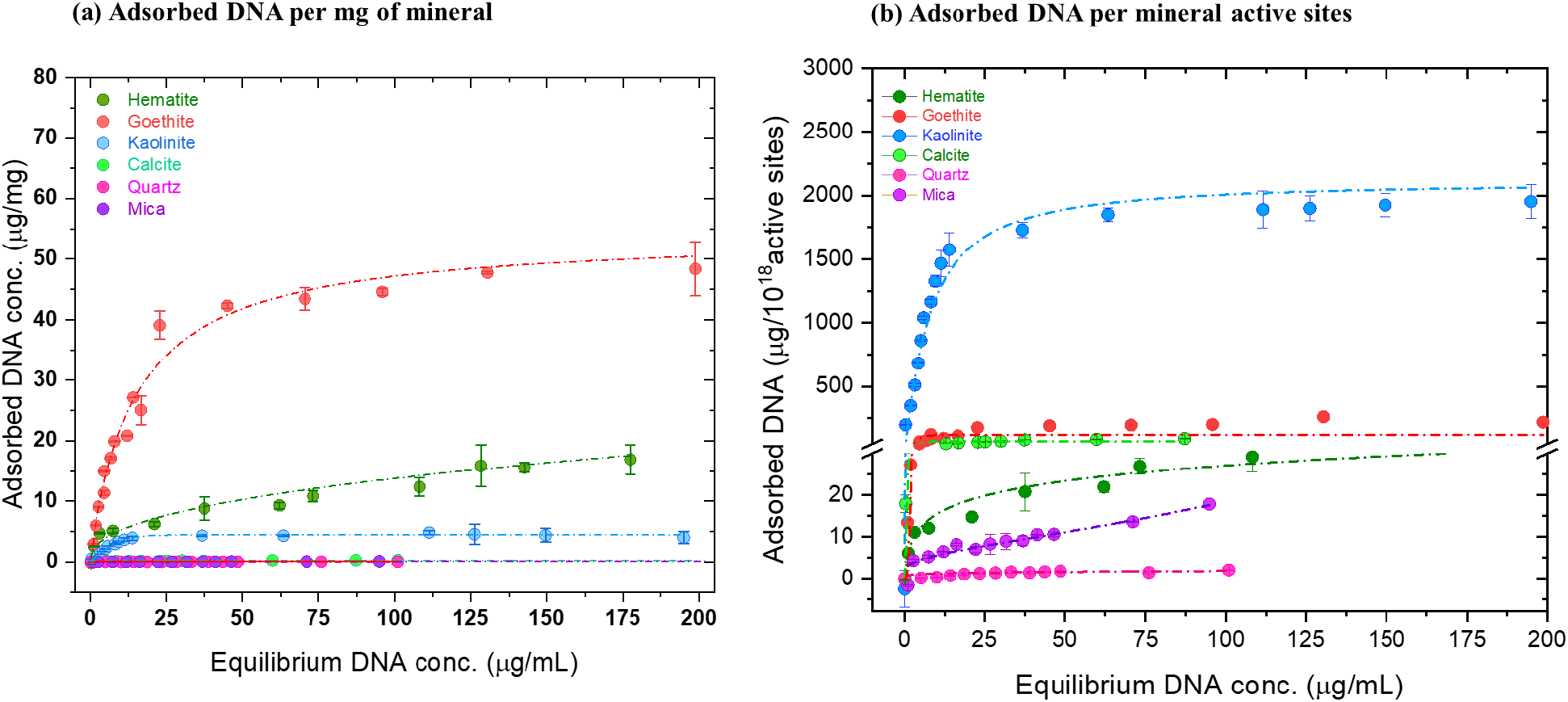
Equilibrium adsorption of salmon sperm DNA in saline under physiological conditions. Serially increasing concentrations of salmon sperm DNA (0-200 μg/mL) were mixed with sonicated mineral suspensions in 150 mM NaCl at pH 7. After incubation for 16 hrs, the amount of adsorbed DNA was estimated using absorbance at 260 nm for the indicated minerals. (a) μg of DNA adsorbed per milligram of mineral and (b) μg of DNA adsorbed per 10^18^ active sites present on each mineral. Langmuir or Freundlich fitting was performed to generate the adsorption isotherms for each DNA-mineral pair and are indicated by the dashed lines. (Langmuir fitting: Goethite, kaolinite, calcite, mica ; Freundlich fitting: Hematite and quartz). The data are representative of three experiments and are plotted as mean ± S.D.

**Figure 2:**
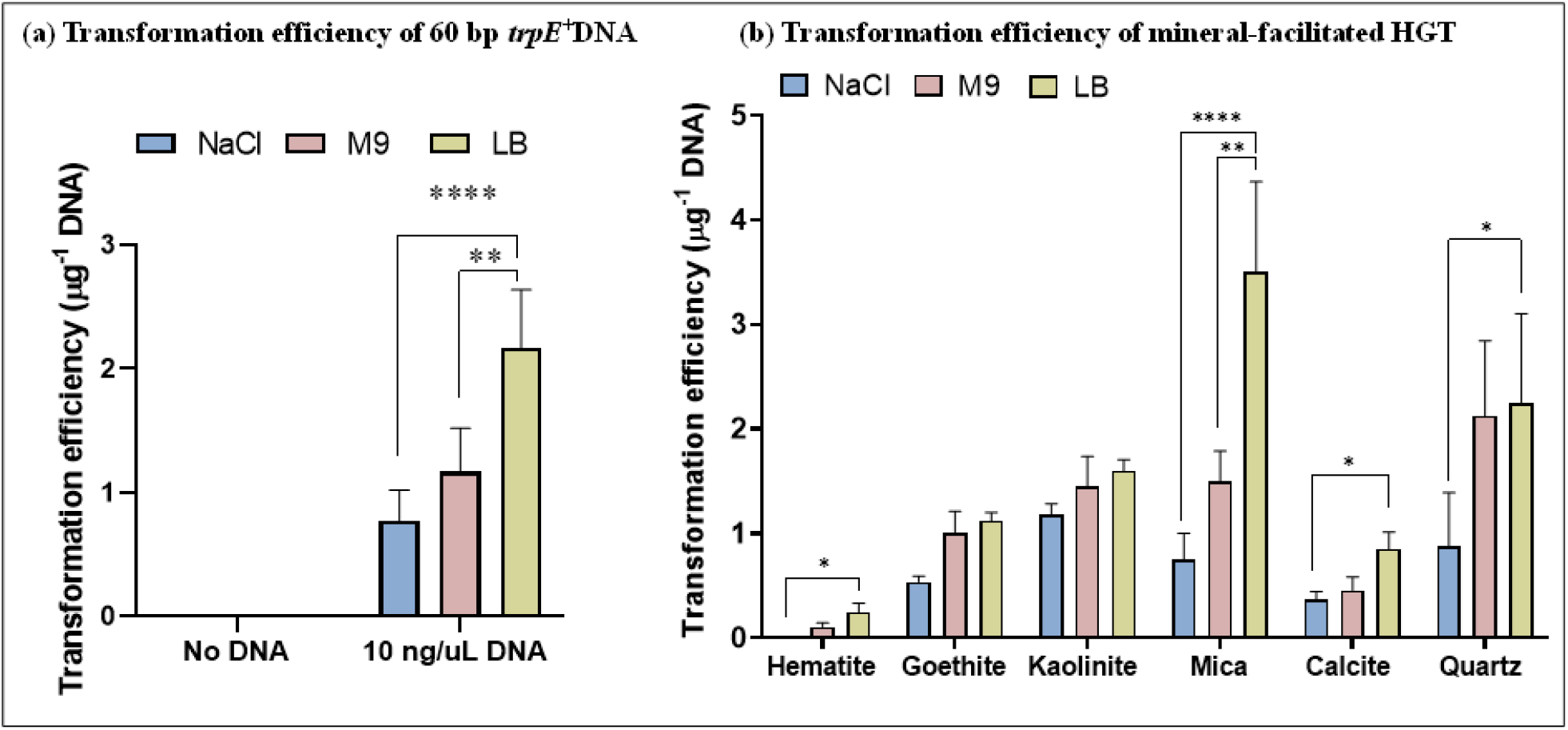
*A. baylyi* is transformed by mineral-adsorbed DNA under nutritionally poor and rich conditions. Competent *trpE*^-^ *A. baylyi* cells were exposed to 60bp *trpE*^*+*^ DNA fragments adsorbed to different minerals as indicated. Recovery of the cells was performed in either saline (150 mM NaCl), M9 supplemented with tryptophan or LB medium for 2 hrs. The transformants were screened using M9 lacking tryptophan and reported as transformation efficiency per μg DNA (number of transformants recovered/μg DNA) obtained using (a) Free DNA and (b) Mineral-adsorbed DNA. The data are representative of three biological replicates and are plotted as mean ± S.D. Statistical analysis was performed using two-way ANOVA.

We also plotted the adsorbed DNA as a function of number of mineral surface active sites and found that when represented as μg DNA/ 10^18^ active sites, kaolinite showed the highest adsorption capacity followed by goethite, calcite, hematite, mica and quartz (Fig. 1(b)). Plotted as μg DNA/ surface area, the order is: goethite, kaolinite, calcite, hematite, mica and quartz (Fig. S1.2). The mass of mineral (mg), mineral surface area (m^2^), DNA concentration (μg/mL), amount of DNA adsorbed (μg/mg) and the derived adsorption parameters in the experiments are listed in Table S2.

#### Transformation

The transformation efficiency of *trpE*^*+*^ DNA into the parent *A. baylyi* strain lacking *trpE* (referred to as *trpE*^-^) was estimated by screening the transformants on M9 agar plates lacking exogenous tryptophan. Cells incubated with 10 ng/μL suspended DNA served as the positive control. For all the nutrient conditions, the transformation efficiency of *A. baylyi* ranged from 0.1 to 4 cells/μg DNA and was strongly dependent on the mineral type (Fig. 2). Bacteria exposed to the negatively charged minerals, quartz and mica, which had the lowest DNA adsorption capacity, displayed the highest DNA uptake efficiency. The positively charged hematite, which had the highest number of active sites, resulted in the lowest transformation efficiency. The transformation efficiencies were generally lowest when cells were grown in saline and highest for M9 and LB media, demonstrating that nutritionally rich conditions enhance DNA uptake owing to more metabolic activity of the cells and more energy allocated for DNA uptake and cell divisions. Moreover, rapidly growing cells have chromosomes with multiple replication forks, offering a high probability and faster uptake frequency of integration of DNA fragments in the bacterial genome. ^23^

To determine if there were any deleterious effects of the minerals which could influence the transformation efficiency, we assessed the cell viability of *trpE*^-^ cells exposed to the minerals. With the exception of goethite and hematite, no significant reduction in the cell viability was observed for the minerals (Fig. S2.1). The growth pattern was consistent when the cells were cultured in either saline (150 mM NaCl), M9 supplemented with tryptophan or LB. Growth was observed only when *trpE*^-^*A*.*baylyi* was plated on M9 agar supplemented with tryptophan, owing to its absolute requirement for tryptophan.

The uptake efficiency of the 60 bp fragments (0.1-4 cells/μg DNA) in our setup is low relative to reported rates of mineral adsorbed plasmids and whole genomes. However, the trend is well in line with previous reports that have shown that transformation frequencies of *Bacillus subtilis, Pseudomonas stutzeri* and *Acinetobacter calcoaceticus* declined with decreasing molecular size of DNA. ^24,25^ The transformation efficiency of *B. subtilis* by sand-adsorbed chromosomal DNA was reported to be 50 times higher when compared to free DNA. ^26^ Other soil bacteria such as *P. stutzeri* and *A. calcoaceticus* showed similar transformation frequencies irrespective of whether the chromosomal DNA was adsorbed to sand or whether it was freely suspended. ^27,28^

#### Importance of interfacial geochemistry for uptake frequency

In general, a direct interaction between a positively charged mineral surface site and a negative phosphate group on the DNA backbone would facilitate a stronger bond than an association to a negatively charged surface site -mediated by the Na^+^ ion as in our case. However, DNA is a large molecule and it has more than one binding site to the mineral surface. The active sites on a mineral surface provide a measure for the density of binding sites and a high site density could lead to a high number of bonds to a DNA molecule. Combined, a high site density, favorable electrostatic interactions and a high surface area would lead to a relatively high adsorption capacity and a DNA molecule strongly bonded to the surface through numerous bonds. The active sites present on the total surface area (**0.025-0.63*10**^**18**^ **sites per 0.06-0.1 m**^**2**^) used for HGT experiments are listed in **Table S1.1**. Hematite has the highest number of total active sites, followed by quartz. Goethite and calcite have the same number of active sites followed by mica. Kaolinite has the least number of active sites per nm^2^ of the mineral. While both hematite (0.64 E18) and quartz (0.5 E18) show a similar number of active sites, in the chemical conditions used in our study, the surface charge for hematite is positive and negative for quartz which will influence the binding strength and adsorption capacity. ^29,30^ Mineral surface charge is important for adsorption dynamics and capacity, but because of the heterogeneity of mineral surface topographic features and site heterogeneity, bulk measures, such as point of zero charge, do not provide the necessary detail to describe how adsorption affinities between mineral and molecules relate to uptake efficiency. Atomic force microscopy provides information on surface features to which DNA associates and fast scanning in liquid gives insight into the mobility of the adsorbed DNA. The mobility is a qualitative measure of the strength of biomolecule bonding to a surface and the availability of adsorption sites.

#### Mineral surface characteristics as drivers for uptake efficiency

We imaged a (001) mica, (001) hematite and (10.4) calcite, in 150 mM NaCl solution. Mica is atomically flat and has a permanent negative charge. The hematite (001) is the most stable hematite face in nature and is used here to represent a general trend for DNA adsorption on iron oxides terraces. The calcite surfaces display charge-dense step edges and terraces that are overall positively charged.

The negatively charged mica surface did not provide a charge landscape in the 150 mM NaCl solution to facilitate a bond strong enough to retain the negatively charged DNA backbone while scanning in liquid. Drying the sample and scanning in ambient air did reveal adsorbed molecules, highlighting that the DNA and the surface had formed a weak association. At the hematite surface, the DNA was readily and stably attached during liquid scanning (Fig. 3 and Fig. S3.1), likely owing to its positive surface charge and high density of active sites. A stack of sequential frames show no-to-minor movement of adsorbed strands (Fig. S3.1(a)). The calcite surface displayed its characteristic step edges and terraces and the DNA molecules aligned along the positively charged step edges. Most of the DNA was associated with the step edge whereas a few DNA molecules were observed on the negatively charged calcite terraces. The DNA adsorbed to the step edges did not move during imaging in liquid conditions, whereas some of the DNA adsorbed to the terraces moved during imaging. A stack of sequential frames shows minor movement of a few selected adsorbed strands (Fig. S3.1(b)). We were not able to track individual strands showing high mobility. The variation in mobility is in line with the variation of defect types on a calcite terrace. ^31^ In light of how the DNA-mineral association influences mobility of the adsorbed DNA, we find the amount of surface binding sites and the bond strength could explain the observed differences in the uptake efficiencies between positively and negatively charged surfaces.

**Figure 3:**
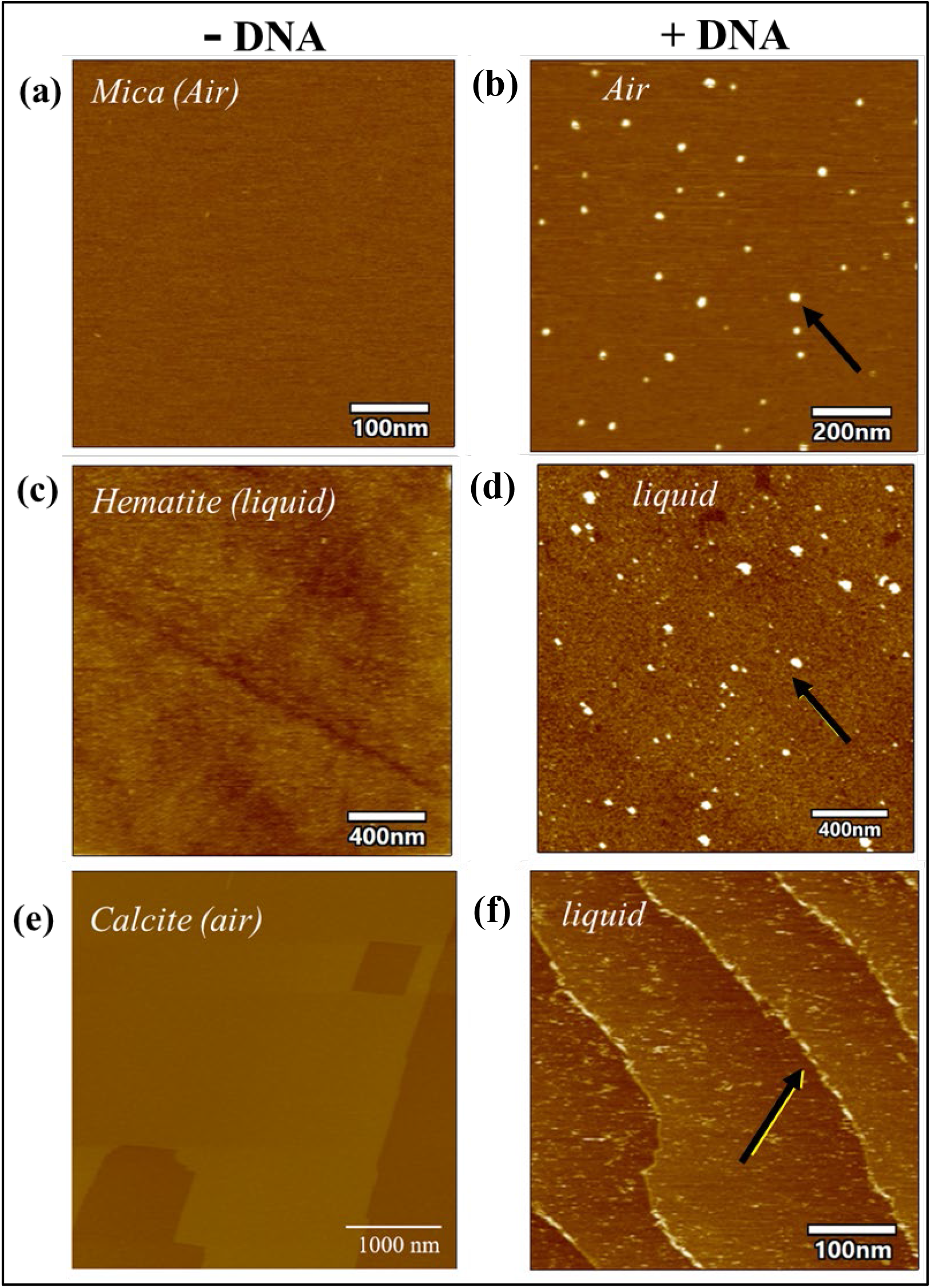
Atomic Force imaging in liquid shows DNA conformation on the mineral surfaces with varying charges. 60bp DNA fragments were allowed to bind to mica (a, b); hematite crystal (c, d) and calcite crystal (e,f) in the presence of 150 mM NaCl at pH 7 and imaged in liquid or air conditions. (Air: Mica; Liquid: Hematite, calcite). Black arrows indicate DNA.

The quartz, mica and kaolinite basal planes will absorb the majority of the DNA through a Na^+^ bridge. The AFM images reveal that the DNA is highly mobile on the mica surface, and it could be argued that the weak adsorption makes the DNA more easily accessible for uptake. The DNA adsorbed on the positively charged clay sites (edge sites and kaolinite octahedral planes) would be strongly bonded and following our logic, available for uptake at a lower rate. The rate at those sites will then be determined by local charge effects and the density of active sites. As for the clay edges and clay octahedral planes, goethite, calcite and hematite can provide a direct electrostatic bond to DNA which is stronger than the bond between DNA and the overall negatively charged minerals. Much of the DNA adsorbed to calcite is adsorbed strongly (in the sense that it is immobilized) which, following our hypothesis, could explain the low uptake frequency. We cannot address though, if the DNA removed by the bacteria was dominantly the small number of mobile strands associated with the terraces or if the strands associated with the terraces in fact represented defect sites. A previous study deriving adhesion forces between a phosphate group and a calcite, and a mica surface does show that calcite bonds to the phosphate group with significantly higher forces than to mica (15 pN vs 1 pN in 100 mM NaCl). ^32^ In that study, the binding force represents an average across terraces and edge sites. Further, Gibbs free energy of binding between phosphate and hematite and mica also confirm that phosphate bonds stronger to Hematite than to mica (5.6 +/-0.9 kt vs. 4.0 +/-0.4 kT). ^33^

For studies of plasmid and chromosomal uptake from minerals into *B*.*subtilis*, differences in uptake efficiencies have also been reported. Competence in *B. subtilis* is known to rely on signaling molecules and it was suggested that mineral adsorption of signaling molecules could decrease competency and hence transformation efficiency.^15,34^ Another potential mechanism explaining the altered transformation rates between minerals is the microbial association with the mineral surface. Strong microbe-mineral interactions have been shown to cause membrane damage,^15,35^ leading to both altered expression of competency genes, and / or reduction of motility. ^34^ We did observe a slight reduction in viability when *A. baylyi* were exposed to goethite and hematite, which may also contribute to the low cell numbers here compared to the other minerals (Fig. S2.1). Untangling these mechanisms is complex and requires knowledge of the mineral surface characteristics such as adsorption capacity, site densities, surface area and charge landscape as well as mapping the involved genes in the bacteria and the bacterial-mineral adhesion landscapes. Common for these possible explanations of variables and limiting factors affecting bacterial DNA uptake from minerals is the association of any of these variables to the mineral, i.e. the DNA –mineral, bacteria-mineral or molecule-mineral associations.

### Significance and scenarios of implications for evolution of bacterial life

Overall, the transformation frequencies are in line with interfacial geochemical principles. Horizontal transformation of fragmented DNA adsorbed to mineral surfaces is influenced by the mineral surface characteristics such that sediments with a high number of active sites and strong bonds would have a high DNA preservation potential, but a lower transformation efficiency. The inverse relationship with preservation and transformation efficiency are in line with our AFM and adsorption data as well and the interfacial arguments presented above. We advocate though, that preservation is more important than the uptake efficiency for long-term evolutionary outcomes and that it is reasonable to assume that each time a cell encounters short and degraded DNA there is a probability that the cell is transformed. First, there is a large amount of fragmented DNA present in the environment, ^20^ second, the incorporation mechanism in the bacterial genome is simple and third, the universality of the mechanism behind the uptake of short and fragmented DNA means DNA integration can, in essence, occur in any cell.

The broad range of settings in which mineral-facilitated HGT can occur may connect organisms through time and space in ways that remain unexplored. On the one hand, bacteria that colonize deposits with historic DNA-mineral complexes can acquire old traits. On the other hand, DNA molecules propagated to distant environments on mineral surfaces may be acquired by organisms far from the DNA-mineral complex where the transported DNA serves as a distal source for new innovations. The age of the DNA would depend on how well the minerals carrying the DNA can preserve (or protect) it from degradation in the given environmental conditions. The remaining question is how microbial evolution is affected by this vast extension of the known temporal and spatial boundaries within which horizontal gene transfer can contribute and how these processes may work and affect natural selection.

Natural selection is a process of adaptation. Adaptation to a novel environment requires heritable variation for selection to act on. Competition and the struggle for existence between individuals will entail that the fittest individuals win the struggle for existence as proposed by Darwin. Since Darwin’s proposal, stochastic mutations that are heritable to offspring are clearly one demonstrated source for variation of the genome. Additionally, horizontal gene transfer is recognized as an important mechanism for the acquisition of beneficial complex traits, and when DNA molecules from one prokaryotic species are taken up by another prokaryote and then inherited by daughter cells, this represents heritable variation. If the genetic material confers a trait and increases the fitness for the offspring, this trait could be selected for. The potential importance of HGT through DNA-mineral complexes lies not only in the movement of traits through time and space. A geological or physical disturbance that creates a novel environment could simultaneously introduce DNA of organisms adapted to those conditions. This proposed mechanism to provide genetic variation differs from where stochastic mutations are invoked. First, the case of mineral-facilitated HGT could influence variation in a non-random fashion since the DNA molecules are specific to the environment. When there is a geological or physical disturbance such as a landslide, a flooding event, or even a new meander on a river, there would be an increased interface between previously established populations and DNA-mineral complexes. Second, the variation and the requirement for adaptation can be linked, in that the process that provides changing conditions that lead to organismal stress and new needs also provides the external DNA molecules. The new needs may arise from the shift in, for example population dynamics (e.g., altered patterns of prey, niches, or friends) or environmental chemistry (e.g., altered localization from oxygen, sulfide, and light). These needs lead to a struggle for existence and, if coupled to new genetic material (via DNA-mineral complexes) conferring a trait that is selected for (increased fitness for the offspring), propelling evolution by natural selection.

The way this proposed mechanism couples stress with increased opportunity is similar to how mutagenesis in cells (prokaryotic, eukaryotic, and cancer cells) can be upregulated in a non-random way by stress. ^36^ For example, stressors such as starvation or drugs induce a DNA-break repair mechanism that is mutagenic. This stress-induced mutagenesis may increase heritable variation and therefore aid genomic solutions (traits) that increase fitness for the offspring of e.g, *E. coli*. In these cases, the mechanism propels evolution in response to new needs via the acquisition of novel and heritable variation; similar to that of mineral-facilitated HGT. Although the coupling of a source for heritable variation (the DNA-mineral complex) and struggle for existence (e.g., as derived by a geological disturbance) as an evolutionary driver is yet less understood as opposed to stochastic mutations, that of mineral-facilitated HGT is not the first suggestion of such mechanism. Consequently, the geological setting can induce and remodel the struggle for existence in a way that by-passes the step of random mutations for increasing fitness. In this scenario, a geologically provided sedimentary and environmental setting for preserving DNA would enhance the adaptive potential of organisms that utilize this route for increased variation.

## CONCLUSION

We demonstrate that mineral-adsorbed 60 bp *trpE*^*+*^ DNA can transform an auxotrophic *A. baylyi* strain with an absolute requirement for exogenous tryptophan into a prototrophic strain that can synthesize tryptophan endogenously. We find that the mineral surface properties have an influence on the uptake frequency such that minerals with a high number of active sites and favorable electrostatics for adsorption show a lower rate of transfer. Considering the ease of incorporation into a genome and the large amounts of DNA associated with sediments, we find that the mixing of modern microbes and ancient DNA in principle represents a pathway for gene transfer, across a scale of time and space, unrivaled by any other avenues. We argue that mineral-facilitated HGT provides a mechanism for rapid adaptations, and we argue that it would provide an efficient path to fitness when there is an abrupt change in the environmental conditions or other evolutionary drivers associated with a shift in the provenance of incoming sediments. Essentially, our results introduce interfacial geochemical and sedimentologic processes as facilitators of evolutionary innovations. Mineral-facilitated HGT could also lead to evolutionary implications that are not yet considered. If the mechanism offers new genomic material at times with increased needs or struggle for existence, it can represent a way to transiently propel evolution in a non-stochastic manner.

## MATERIALS AND METHODS

### Minerals and their characterisation

The Hematite {00.1} face used for AFM was a polished, single crystal of hematite (HEM). The face was cleaned in 1 M NaOH at 60 °C for 1 h to remove fatty contamination. Subsequently it was sonicated for 20 minutes, rinsed with molecular grade water and UV-ozone cleaned for 20 min. Goethite was synthesized by modifying the protocol of Martina *et al*., 2022.^41^ Briefly, 24 grams of Ferric (III) nitrate nonahydrate (Fe(NO_3_)_3_.9H_2_O) (Sigma) were added to 300 mL MilliQ water while stirring vigorously using a magnetic stirrer. The solution was simultaneously titrated with 2 M NaOH until it reached a pH of 9. The solution was stirred for another 30 mins to allow the reaction to reach completion. The resulting brown precipitate was allowed to age to the yellow-coloured goethite by incubation at 60°C for 24 hrs. Goethite was then washed thoroughly with MilliQ water until no nitrate was detected in the supernatant (Nitrate detection strips, Supelco, Sigma). The purity of the synthesized goethite was confirmed using FTIR spectroscopy. Kaolinite (Kga-1b) was used a purchased from the Clay Minerals Society.

Calcite were purchased from Sigma and cleaned according to Belova et al. ^37^ Quartz was purchased as large crystals which were ground and sieved to obtain <50 μm particles. The Mica powder was from an outcrop in the French alps and the mica substrate used for AFM is V1 grade purchased at Ted Pella Inc.

### Adsorption isotherms of DNA to minerals

The minerals were weighed according to their surface area as listed in Table S1.1. Each mineral was disaggregated using a sonicator for 30 mins and autoclaved at 121°C, 15 lbs pressure for 20 minutes to remove any microbial contamination and resuspended in 1 mL sterile 150 mM NaCl at pH 7. The mineral suspensions were mixed with 0-200 ng/μL salmon sperm DNA in triplicates (also at pH 7) (Sigma). The mineral-DNA suspensions were incubated at room temperature for 16 hrs under constant shaking at 70 rpm. Post incubation, the suspensions were centrifuged at 14000 rpm for 5 mins and the DNA released in the supernatant was measured using absorbance at 260 nm using a UV-visible spectrophotometer (Eppendorf). The amount of mineral-adsorbed DNA was estimated using the following equation:

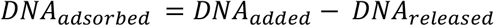

Adsorption isotherms were plotted as equilibrium DNA concentration on the x axis and adsorbed DNA concentration on the y axis. The data were fitted using either the Langmuir equation or the Freundlich equation. **The Langmuir model for adsorption followed the equation:**

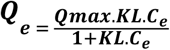

Where ***Q***_e_ is the amount of DNA adsorbed per unit mass of mineral

***K***_***L***_ is the Langmuir adsorption constant

***Qmax*** is the maximum adsorption capacity of the mineral

***C***_e_ is the equilibrium concentration of the DNA

**The Freundlich adsorption isotherm follows the equation:**

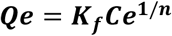

where ***Qe*** is the amount of DNA adsorbed per unit mass of mineral

***Ce*** is the equilibrium solution concentration of the DNA,

***K***_***f***_ is the Freundlich adsorption constant, ***n*** is the empirical constant.

### Atomic Force Microscopy and sample preparation for liquid and air imaging

AFM imaging in both air and liquid was performed using a Cypher VRS system (Oxford Instruments). Imaging in air was performed using Tap150 tip with spring constants between 1.5 and 15 N/m and a resonant frequency of 150 kHz. For liquid imaging, AC40 tip (Olympus) (Spring constant 0.1 N/m; Resonant Frequency 500 kHz). A single-phase crystal of hematite was used for AFM imaging. 60 bp ds DNA at a final concentration 0.5 ng/μL in 150 mM NaCl was added to the surface of the hematite crystal and imaged under liquid mode. A clean and uniform calcite surface was obtained by cutting the calcite crystal using a scalpel blade and immediately removing the dust using a gentle N_2_ stream. Mica was cleaved using the tape-stripping method to obtain a clean surface. Scan angle, scan rate, and set-point were systematically varied and multiple sites on every sample were imaged.

### Bacterial strains, culture conditions and competent cell preparation

*Acinetobacter baylyi* BD413 was kindly provided by Klaus Harm, University of Tromsø, Norway and carries the *trpE27* mutation making the strain auxotrophic for tryptophan. TrpE codes for anthranilate synthase subunit I which a part of the tryptophan biosynthesis pathway. G to A transition at the 883^rd^ position of the 1494 bp long *trpE* ORF results in an amino acid change (E295K) making the gene product nonfunctional ^21^. To prepare competent cells, fresh overnight culture was diluted 1:100 in LB and incubated at 30°C under shaking at 180 rpm till an O.D of 0.3 was obtained (∼4 hrs), corresponding to ∼10^7^ cells. The cells were centrifuged at 5000 rpm at 4°C for 10 mins and resuspended in prechilled LB containing 20% glycerol, aliquoted and stored at -80°C until further use.

### Horizontal gene transfer experiments in the presence of mineral-DNA complexes

For HGT experiments, the DNA used was a 60 bp double stranded DNA fragment with the sequence (3’CAATACCATCTTCCAAGCGTGACAGGATTTCGGGTGATGAGCCAACAATGTGAAAAGGCG 5’ 5’GTTATGGTAGAAGGTTCGCACTGTCCTAAAGCCCACTACTCGGTTGTTACACTTTTCCGC-3’) procured from Integrated DNA Technologies (IDT, Belgium). Autoclaved and sonicated minerals were mixed with the appropriate concentration of DNA (as derived from the adsorption isotherm data, see Table S1.2). The brief experimental strategy is depicted in Figure MM-1 in this section.

**Figure MM-1:**
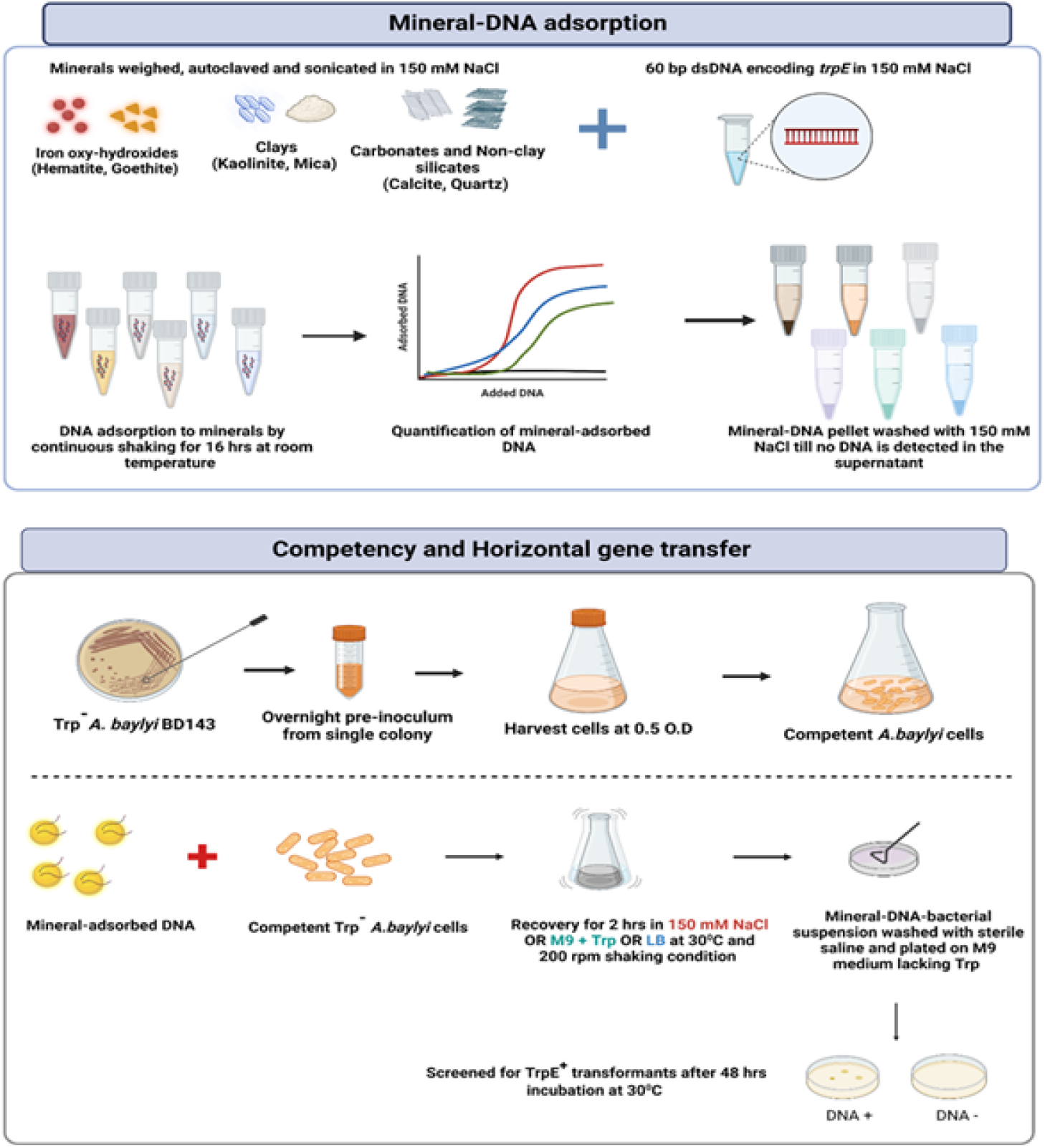
Schematic outlining the experimental strategy used for preparation of mineral-DNA complexes for adsorption as well as competent cell preparation and mineral-facilitated horizontal gene transfer.

We aimed to add a concentration of DNA to the transformation experiments where the mineral was not fully saturated with DNA and the adsorption isotherms were used as guides to select the DNA:mineral ratio applied in the transformation experiments.

The initial DNA concentration was chosen such that the DNA adsorbed by the positively charged minerals (hematite, goethite, kaolinite and calcite) was ∼ 10 ng/μL whereas for the negatively charged minerals (mica and quartz), it was ∼ 1 ng/μL. The mineral-DNA suspensions were shaken for 16 hrs at room temperature at 70 rpm and centrifuged at 5000 rpm for 30 mins. The residues were washed with 1 mL sterile saline until no free DNA was detected in the supernatant (estimated by measuring absorbance at 260 nm). The mineral-DNA complexes were then used for transformation experiments. For horizontal gene transfer experiments, competent cells were thawed on ice and washed once with sterile saline. The cells were then added to the mineral-DNA complex, resuspended in 150 mM NaCl or M9 supplemented with tryptophan or LB and incubated for 2 hrs at 30°C under shaking at 100 rpm. Post incubation, the mineral-DNA-bacteria complexes were washed once and resuspended in 300 μL of sterile saline. The cells were plated on M9 agar containing no tryptophan and incubated at 30°C for 48 hrs to screen for transformants. As a control experiment, *A. baylyi* cells were exposed to the minerals in the absence of DNA and plated both on M9 supplemented with tryptophan as well as without tryptophan. The transformation efficiency per μg DNA was calculated as follows:

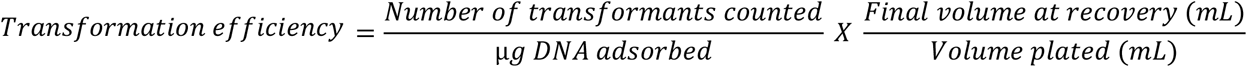

### Statistical analysis

All experiments were performed in triplicates and are plotted as mean ± S.D. Statistical analysis was performed using two-way ANOVA using GraphPad Prism 8 where * represents p<0.05; ** p<0.01; *** p<0.001 and **** p<0.0001.

### Atomic force microscopy

#### Samples and relevance

AFM is limited to flat surfaces which makes minerals such as mica and calcite suitable. Considering overall surface charge DNA is expected to display a similar adsorption behavior between mica and quartz. The basal planes of kaolinite are similar to the structure of mica and hence we mica was chosen to represent these minerals. Calcite surfaces display charge dense step edges and terraces that are overall positively charged. The iron oxides we applied (goethite and hematite) are nanoparticulate and display a range of mineral surfaces and local charge densities. It is not feasible to image the individual faces with AFM and we applied a flat 001 hematite surface to represent a general trend for DNA adsorption on iron oxides terraces. DNA adsorption to step edges on iron oxides are expected to behave similar to DNA adsorbed on calcite step edges.

#### Sample preparation, imaging and analysis

Mica sheet was stuck on the AFM stub using glue and was cleaved using the tape stripping method to obtain a clean sterile surface. For hematite, a flat {00.1} crystal was used which was cleaned with 1 M NaOH and sterilised under UV-ozone before use. A smooth surface for calcite was obtained by cutting a calcite block with a scalpel. All surfaces were cleaned with a gentle stream of Nitrogen gas.

A stock DNA concentration of 1 ng/uL in 150 mM NaCl was used for all the AFM experiments. DNA was diluted to 0.5 ng/uL using 150 mM NaCl before being added to the mineral surfaces (hematite, mica and calcite). For liquid imaging, the mineral surface was covered with ∼15-20 uL 150 mM NaCl and imaged directly under AFM. Thereafter, DNA was introduced, allowed to incubate for 2 mins and imaged.

For imaging under air conditions, 0.5 ng/uL DNA was added to the mineral surface and allowed to incubate for 2 mins. Post incubation, the mineral surface was gently rinsed with 2X filtered MilliQ water, blotted with a kimwipe and dried completely under a gentle stream of N_2_ gas.

For tracking the movement of DNA across hematite and calcite, images in the TIFF format were retrieved from AFM movies using software Igor Pro v 6.38 and processed using MATLAB. In MATLAB, hematite images pixel values were converted into double data type and then the negative images were obtained by subtracting the scaled pixel values to 1. After this step, the background became light gray and the particles on the surface appeared as black dots. A mask was applied to remove most of the particles (black dots) in the images. The mask assigned the average background pixel value to all the pixels with a value higher than 0.89 except in 5 specific locations in the images (those corresponding to the particles of interest). Calcite images were also processed in MATLAB and their pixel values were converted into double data type. In this case, instead of obtaining the negative image, the background features were highlighted by using a mask. This mask assigned a value of 0 to all the pixels that had values higher than a number in the range from 0.75 to 0.9. The selection of a value within this range depended on the overall pixel values in the image: images with lower pixel values required a lower value for the mask and images with higher pixel values required a higher value. After this, the processed images were exported in TIFF format and superimposed using the background features as guides in PowerPoint. The DNA molecules in the images (highlighted as black dots) were tracked throughout the images and their area of movement was coloured green.

## AUTHOR CONTRIBUTIONS

TV and KKS designed the study and wrote the manuscript, TV carried out the bacterial experiments supported by SH. TV and KKS planned and carried out the AFM imaging. CC carried out image stacking. SA and EH contributed with evolutionary context. MB was deeply involved in designing and planning bacterial experiments. All authors contributed to writing the manuscript.

## ACKNOWLEDGEMENTS

The authors would like to thank Stanislav Jelavić for discussions on DNA-mineral affinities. This work was supported by a research grant from VILLUM FONDEN (00025352).

