## Supporting Information for "Recurrence and propagation of past functions through mineral facilitated horizontal gene transfer"

**Supporting section 1: minerals**

1.1: Adsorption and mineral data

**Table S1.1: Properties of the minerals used in the study.**

| **Mineral** | **Type** | **Specific surface area (SSA) m^2^ g^-1^** | **Surface area of the minerals used for HGT (m^2^)** | **Active sites per nm^2 (as obtained from references)^** | **Active sites on the total surface area used for HGT (nm^2^)** |
| --- | --- | --- | --- | --- | --- |
| **Hematite** | Iron oxy-hydroxide | 108.5 | 0.1 | 6.36 ^38^ | 0.636*10^18^ |
| **Goethite** |  | 90.1 | 0.06 | 5.5 ^39^ | 0.33*10^18^ |
| **Kaolinite** | Clay minerals | 11.7 | 0.1 | 0.25 ^40^ | 0.025*10^18^ |
| **Mica** |  | 1.5 | 0.06 | 5 ^41^ | 0.3*10^18^ |
| **Calcite** | Carbonate | 0.31 | 0.06 | 8.5 ^42^ | 0.51*10^18^ |
| **Quartz** | Non-clay silicate | 1.5 | 0.06 | 4 ^43^ | 0.24*10^18^ |

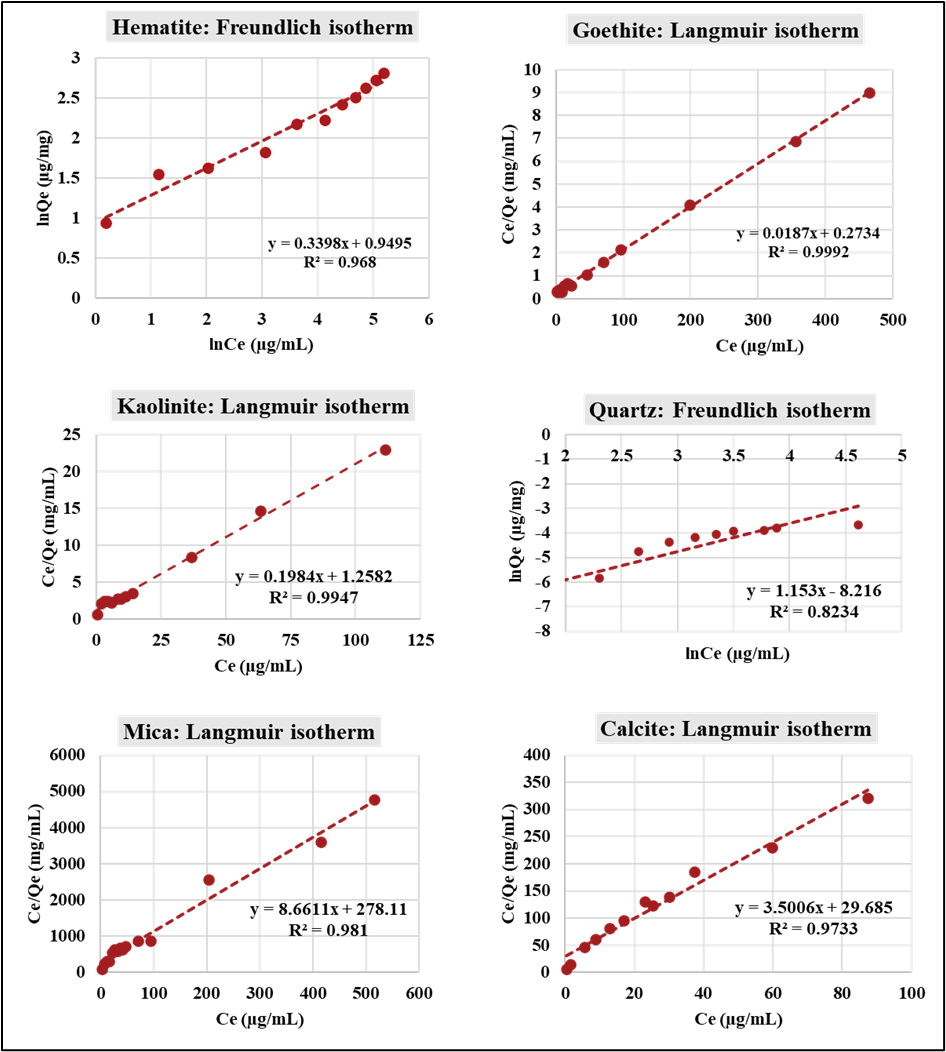
**Figure S1.1: Adsorption isotherm plots for DNA binding on minerals.** Adsorption data for salmon sperm DNA and the different minerals were used to fit either the linear Langmuir equation for goethite, kaolinite, mica and calcite or the linear Freundlich equation for hematite and quartz. Red dots represent the adsorption data points and the dotted line refers to Langmuir or Freundlich fit. ‘Ce’ refers to the equilibrium DNA concentration and ‘Qe’ is the adsorbed DNA concentration. The data are representative of three experiments and are plotted as mean.

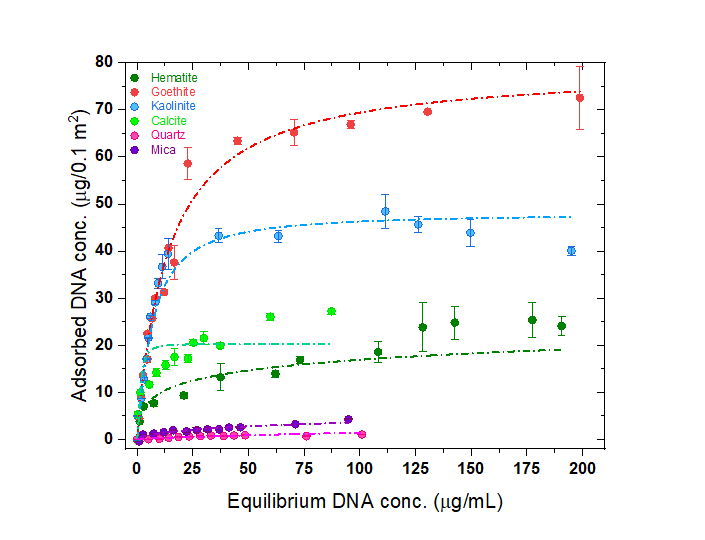

**Figure S1.2: Equilibrium adsorption of salmon sperm DNA corresponding to 0.1 m^2^ of the mineral surface area in saline under physiological conditions .** Serially increasing concentrations of salmon sperm DNA (0-200 µg/mL) were mixed with sonicated mineral suspensions in 150 mM NaCl at pH 7. After incubation for 16 hrs, the amount of adsorbed DNA was estimated using absorbance at 260 nm for the indicated minerals. Langmuir or Freundlich fitting was performed to generate the adsorption isotherms for each DNA-mineral pair. (Langmuir fitting: Goethite, kaolinite, calcite, mica ; Freundlich fitting: Hematite and quartz). The data are representative of three experiments and are plotted as mean ±S.D.

1.2 Information on mineral morphologies and topographies

Hematite and goethite are nano rods composed of several crystal faces such as (100), (110) and (021) with slightly distinct charges. In general, goethite displays a higher density of hydroxyl groups and positive charges than hematite. Mica and kaolinite are both clay minerals and have permanent negatively charged basal planes and positively charged edge sites. In contrast to mica, kaolinite has two types of basal planes where one is composed of Al-octahedra, which are slightly positive in our experiments. The basal planes are atomically flat. Calcite is overall positively charged, and the major crystal face exposed is the (10.4) face. The local topography on the calcite surface is composed of atomically flat terraces and the surface contains obtuse and acute step edges which carry a stronger positively charge density than the terraces.

**Table S1.2: Mineral weights and DNA concentrations used in the study.**

| **Mineral type** | **Weight of the mineral (mg)** | **DNA added**  **(µg µL^-1^)** | **DNA adsorbed**  **( µg mineral wt^-1^)** | **Adsorption parameters** | | |
| --- | --- | --- | --- | --- | --- | --- |
|  |  |  |  | **Langmuir adsorption** | | |
|  |  |  |  | **Q_max_** | **K_L_** | **r^2^** |
| Goethite | 0.7 | 0.02 | 11±2.6 | 53.5 | 0.07 | 0.99 |
| Kaolinite | 10 | 0.015 | 11±2 | 5.04 | 0.16 | 0.99 |
| Mica | 40 | 0.075 | 1.3±0.6 | 0.11 | 0.03 | 0.98 |
| Calcite | 200 | 0.015 | 9.3±1.6 | 0.3 | 0.12 | 0.97 |
|  |  |  |  | **Freundlich adsorption** | | |
|  |  |  |  | **K_F_** | **n** | **r^2^** |
| Hematite | 1.5 | 0.03 | 8.3±1.1 | 2.58 | 2.94 | 0.97 |
| Quartz | 40 | 0.05 | 1.3±0.6 | 3700 | 0.86 | 0.82 |

Q_max_: maximum adsorption capacity (µg/mg), K_L_: Langmuir constant, K_F_: Freundlich constant, n: Freundlich exponent

**Supporting section 2: Cell viability**

To study the effects of minerals on bacterial viability, we exposed *A. baylyi* to fixed mineral concentrations (see Table S1.2 for concentrations) in saline, M9 supplemented with tryptophan or LB for a period of 2 hrs under shaking conditions. The mineral-bacterial suspensions were then plated on M9 supplemented with tryptophan, incubated at 30^0^C for 48 hrs and the colonies were counted. Except for goethite and hematite, no significant reduction in the cell viability was observed for the minerals.

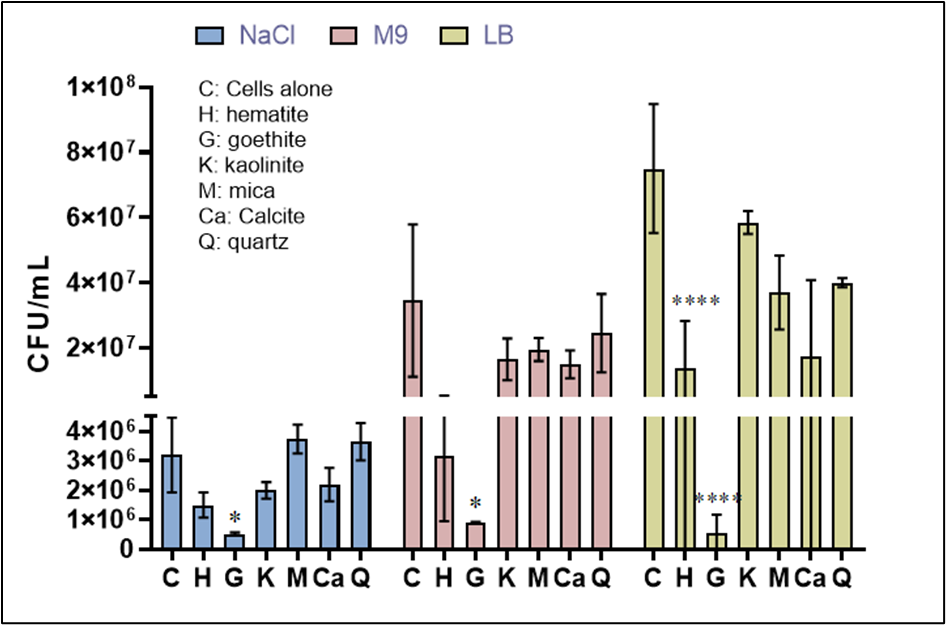

**Figure S2.1: Effect of minerals on cell viability of *A.baylyi*.** Competent *A. baylyi* cells were exposed to the various minerals as indicated for a period of 2 hrs in either saline (150 mM NaCl), M9 supplemented with tryptophan or LB medium. Post exposure, the cells were plated onto M9 supplemented with 50 mg/L tryptophan and incubated at 30^O^C for 48 hrs. Colonies were counted and reported as CFU/mL. The data are representative of three biological replicates and are plotted as mean ± S.D. For statistical analysis, two-way ANOVA was performed for all minerals with respect to cells alone (For eg.: Cells alone vs cells in hematite grown in NaCl.)

**Supporting section 3. AFM**

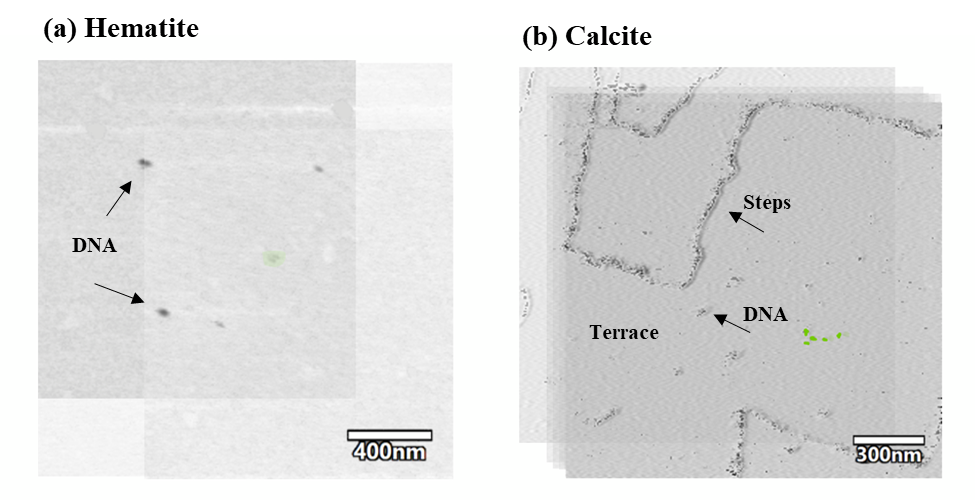

**Figure S3.1: Time-dependent tracking of DNA using AFM image stacks of hematite and calcite.** (a) Overlay of 6 hematite images taken during a time span of 6 minutes while tracking 5 DNA molecules (DNA molecules are highlighted as black dots); (b) Overlay of 6 calcite images taken during a time span of 6 minutes while tracking 5 individual DNA molecules anchored to a terrace.

**Images in the TIFF format** were retrieved **from AFM movies using software Igor Pro v 6.38 and** processed using MATLAB. In MATLAB, hematite images pixel values were converted into double data type and then the negative images were obtained by subtracting the scaled pixel values to 1. After this step, the background becomes light gray and the particles on the surface appear as black dots. A mask was applied to remove most of the particles (black dots) in the images. The mask assigned the average background pixel value to all the pixels with a value higher than 0.89 except in 5 specific locations in the images (those corresponding to the particles of interest). Calcite images were also processed in MATLAB and their pixel values were converted into double data type. In this case, instead of obtaining the negative image, the background features were highlighted by using a mask. This mask assigned a value of 0 to all the pixels that had values higher than a number in the range from 0.75 to 0.9. The selection of a value within this range depended on the overall pixel values in the image: images with lower pixel values required a lower value for the mask and images with higher pixel values required a higher value. After this, the processed images were exported in TIFF format and superimposed using the background features as guides in PowerPoint. The DNA molecules in the images (highlighted as black dots) were tracked throughout the images and their area of movement was coloured green.
